# Comparative analysis of the caecal tonsil transcriptome in two hen lines experimentally infected with *Salmonella* Enteritidis

**DOI:** 10.1101/2022.06.03.494662

**Authors:** Anaïs Cazals, Andrea Rau, Jordi Estellé, Nicolas Bruneau, Jean-Luc Coville, Pierrette Menanteau, Marie-Noëlle Rossignol, Deborah Jardet, Claudia Bevilacqua, Bertrand Bed’hom, Philippe Velge, Fanny Calenge

## Abstract

Managing *Salmonella enterica* Enteritidis (SE) carriage in chicken is necessary to ensure human food safety and enhance chicken breeding viability. *Salmonella* can contaminate poultry products, causing human foodborne disease and economic losses for farmers. Both genetic selection for a decreased carriage and gut microbiota modulation strategies could reduce *Salmonella* propagation in farms.

Two-hundred and twenty animals from the White Leghorn inbred lines N and 6_1_ were raised together on floor, infected by SE at 7 days of age, transferred into isolators to prevent oro-fecal recontamination and euthanized at 19 days. Caecal content DNA was used to measure individual *Salmonella* counts (ISC) by droplet digital PCR. A RNA sequencing approach was used to measure gene expression levels in caecal tonsils after infection of 48 chicks with low or high ISC.

The analysis between lines identified 7516 differentially expressed genes (DEGs) corresponding to 62 enriched Gene Ontology (GO) Biological Processes (BP) terms. A comparison between low and high carriers allowed us to identify 97 DEGs and 23 enriched GO BP terms within line 6_1_, and 1034 DEGs and 288 enriched GO BP terms within line N. Among these genes, we identified several candidate genes based on their putative functions, including *FUT2* or *MUC4*, which could be involved in the control of SE infection, maybe through interactions with commensal bacteria. Altogether, we were able to identify several genes and pathways associated with differences in SE carriage level. These results are discussed in relation to individual caecal microbiota compositions, obtained for the same animals in a previous study, which may interact with host gene expression levels for the control of the caecal SE load.

## Introduction

*Salmonella* is a zoonotic pathogen that can cause human foodborne disease. In 2019, more than 80,000 human salmonellosis cases were confirmed in Europe with 140 reported deaths [1]. Among these cases, more than 9000 were associated with 926 food-borne outbreaks (FBOs), with a large majority (72.4%) caused by the serovar *S. Enterica* Enteritidis (SE). Eggs produced by infected layer hens seem to be the major food vehicle, representing more than 37% of the FBOs. In parallel, in spite of strict hygiene control in farms, the systematic detection of *Salmonella* serovars, and the use of vaccination, the prevalence of *Salmonella* in laying hen flocks increased from 2.07% in 2014 to 3.44% in 2019 [1]. In chicken, the carriage is asymptomatic. The bacteria can persist a long time in the gut and can quickly spread within a contaminated farm via oro-fecal recontaminations between birds [2]. Understanding the impact of factors such as host genetics or gut microbiota on *Salmonella* carriage, and even more their combined impact, could lead to innovative strategies to reduce *Salmonella* transmission and ensure human food safety.

The caecal tonsil is a major barrier controlling the entry of bacteria in the organism [3,4] and is therefore a tissue particularly relevant for identifying host factors potentially involved in the control of SE. Several studies have been conducted on the caecal tonsil transcriptome. In particular, they have helped to identify biological processes associated with resistance to *S*. Enteritidis [5–7], *S*. Typhimurium [8,9] and *S*. Pullorum [10]. Nevertheless, in these studies, the impact of host genetics on gene expression was not examined. The expression of specific immune genes has been compared between the two experimental inbred chicken lines 6_1_ and 15I, but not whole transcriptome [11]. The impact of host genetic variations was considered in a recent study of the caecal tissue transcriptome after *Campylobacter* colonisation. Comparisons between the experimental White Leghorn inbred chicken lines 6_1_ and N led to the identification of a large number of differentially expressed genes, which may underlie variation in heritable resistance to the pathogen [12].

The host genetic background is an important factor for the outcome of *Salmonella* infection in chicken. A number of quantitative trait loci (QTL) and candidate genes associated with *Salmonella* resistance have been identified [13]. In the inbred lines N (resistant) and 6_1_ (susceptible) in particular, several QTLs with low to moderate effects were identified [14–16]. However, no causal gene could be pinpointed due to the large size of the QTL genomic regions. More generally, only a few genes have been identified for their direct implication in the control of SE load in chicken, and knowledge is lacking about the mechanisms leading to genetic resistance. For the studies conducted on *Salmonella* carriage in the N and 6_1_ lines, birds were reared together on floor after infection, thus allowing *Salmonella* orofecal recontamination between birds. In the present study, we used another infection model, making use of isolators. Previously tested on the experimental White Leghorn line PA12, this model showed a strong reduction of oro-fecal recontaminations, leading to much increased *Salmonella* individual variation among birds [2]. It is therefore an interesting model to identify birds with highly contrasted carriage levels, in order to facilitate the identification of host genes involved in these differences.

Another major protective barrier against *Salmonella* infection is the host gut microbiota [17,18]. The protective role of the resident gut microbiota toward SE colonisation has been established for a long time [19] and has been called “competitive exclusion” [20,21]. In several studies, the analysis of the impact of *Salmonella* infection on the host microbiota composition led to the identification of specific gut bacteria and biological mechanisms which could prevent *Salmonella* colonisation in chicken [22,23]. Moreover, specific gut microbiota compositions determine the super and low-shedder phenotypes in the model hen line PA12 [24]. Besides, more generally the existence of a genetic control of the microbiota composition in chicken has been demonstrated using several genetic backgrounds [25–27]. QTLs and single nucleotide polymorphisms (SNPs) associated with the abundance of bacteria have been identified [28,29]. Given the importance of the gut microbiota composition for resistance to SE carriage, we hypothesize that one of the mechanisms by which host genetics acts on the SE carriage level may be through an effect on the gut microbiota composition. This hypothesis was not contradicted by our latest study, in which we identified significant differences in microbiota composition between line N and line 6_1_, and between low and high *Salmonella* carriers in line 6_1_. We were able to identify some bacteria associated with *Salmonella* resistance, such as the *Christensenellaceae* family [30].

In the current study, we performed an integrative analysis using information about caecal gene expression and caecal microbiota composition in two distinct genetic lines: the inbred chicken lines N and 6_1_, respectively resistant and susceptible to SE infection. The objectives of this study were to:

i. identify differentially expressed genes between genetic lines in the caecal tonsils after SE infection, in order to identify potential pathways involved in the genetic resistance to SE;
ii. identify genes and pathways associated with SE resistance within line (low vs high carriers);
iii. link the results to those we obtained on the caecal microbiota [30] to identify potential interactions between the host and its microbiota that could explain differences in resistance to SE infection.

## Results

Two-hundred and forty animals from the two experimental White Leghorn inbred lines N and 6_1_ were raised together on floor until 7 days of age. Then, chicks were challenged with *Salmonella* enterica Enteritidis (SE) LA5 by oral infection and separated into four isolators. Two independent replicates (n=120) were conducted with a total number of 240 chicks. No clinical signs of disease were observed on the animals. Caecal contents and caecal tonsils were collected at 12 days post infection.

The abundance of SE in caecal contents was measured by ddPCR and, as described previously, significant differences of *Salmonella* abundance were observed between lines for the two experiments [30]. The observed variability of carriage allowed us to identify extreme low and high carriers within each line, and 48 extreme animals were selected for the caecal tonsil RNA extraction, balancing the “experiment”, “isolator” and “sex” factors. Different groups were defined as described in Fig 1 according to the line, class and experiment factors. Means, standard deviations and p-values according to these groups are also given in Fig 1.

**Fig 1.**
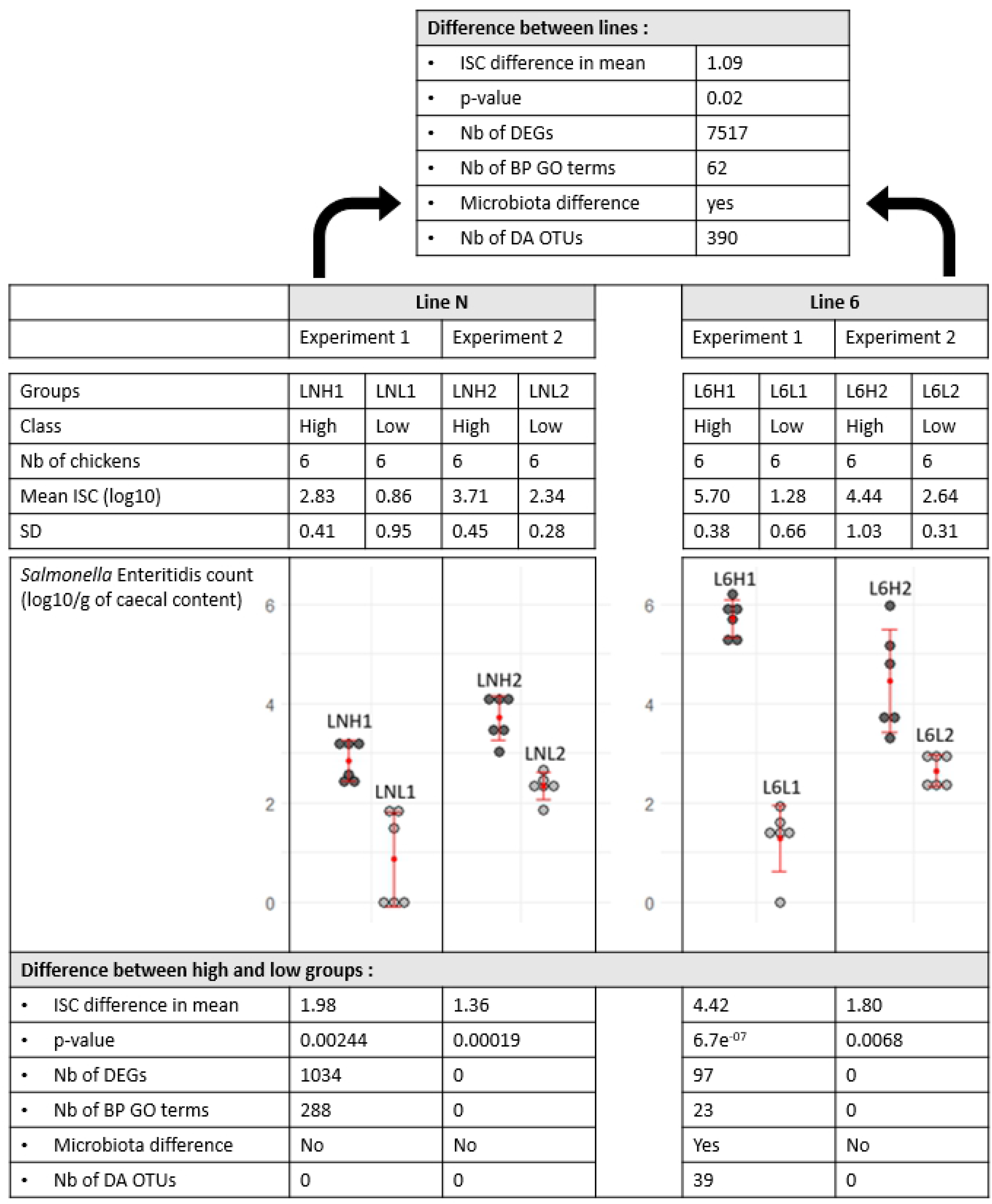
Summary of the results of comparisons between lines and between low and high carrier classes within lines according to the experiment. Mean (red points) and standard deviation (SD; red bars) of the *Salmonella* Enteritidis abundance at 12 dpi in caecal contents (log10/g of caceal contents) of chicken groups infected with SE according to the line and experiment. Difference in mean between low and high carriers according to the line and experiment and t-test p-value. Difference in mean between lines and t-test p-value. Results of the DEG and BP GO term enrichment analyses. Results of the 16S analysis from the study [30].

### Differentially expressed genes (DEGs) in caecal tonsils between lines

On average, more than 40M reads were sequenced for each of the 48 samples. After quality control, a total of 24,356 expressed genes were identified and used for the following analyses. Using all 48 samples, a principal component analysis (PCA) showed a distinct clustering between lines (Fig 2). Two ANOVA analyses on the PCA1 and PCA2 components showed that gene expression is significantly affected by line and sex (S1 File). A differential analysis with DESeq2 allowed the identification of 7,516 DEGs between the two lines (p-adj<0.05) among which 3,944 were up- and 3,572 were down-regulated (S1 Fig. and S1 Table).

**Fig 2.**
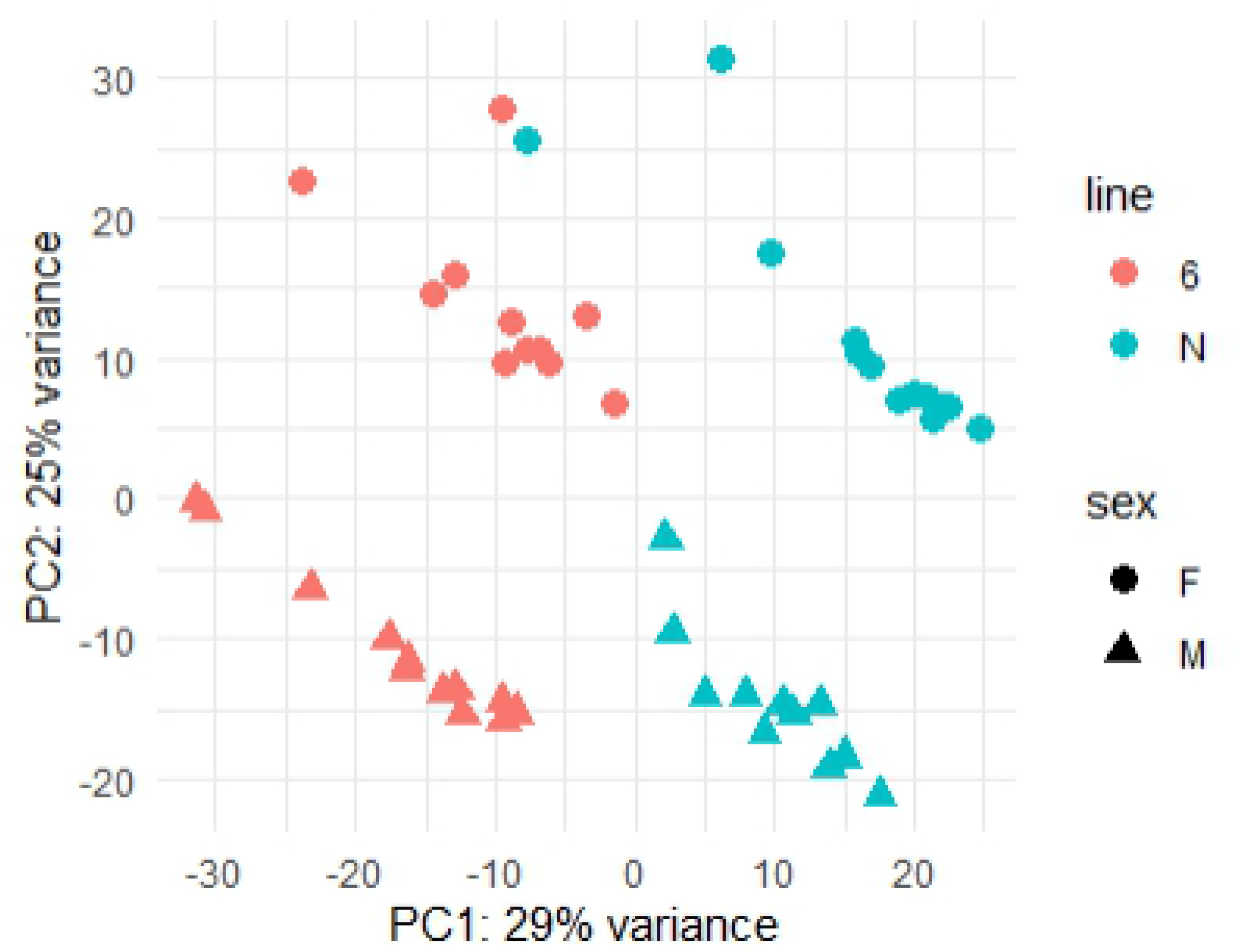
Principal component analysis of the gene expression in caecal tonsils in all samples. The first principal component contains 29% of the variance and may be attributed to the line effect and the second principal component contains 25% of the variance and may be attributed to the sex effect.

### DEGs in caecal tonsils between low and high carriers within lines and experiments

Gene expression levels between low and high *Salmonella* carriers within each line and each experiment (L6L1/L6H1, L6L2/L6H2 and LNL1/LNH1, LNL2/LNH2) were compared. A PCA showed a distinct clustering between the high and lower carriers within each of the N and 6_1_ lines in experiment 1 (Fig 3; LNH1 vs LNL1 and L6H1 vs L6L1), but not in the equivalent groups of experiment 2. ANOVA analysis on the PCA1 and PCA2 components showed that gene expression is significantly (P<0.05) affected by low/high classes and by sex in experiment 1, but not in experiment 2 (S1 File).

**Fig 3.**
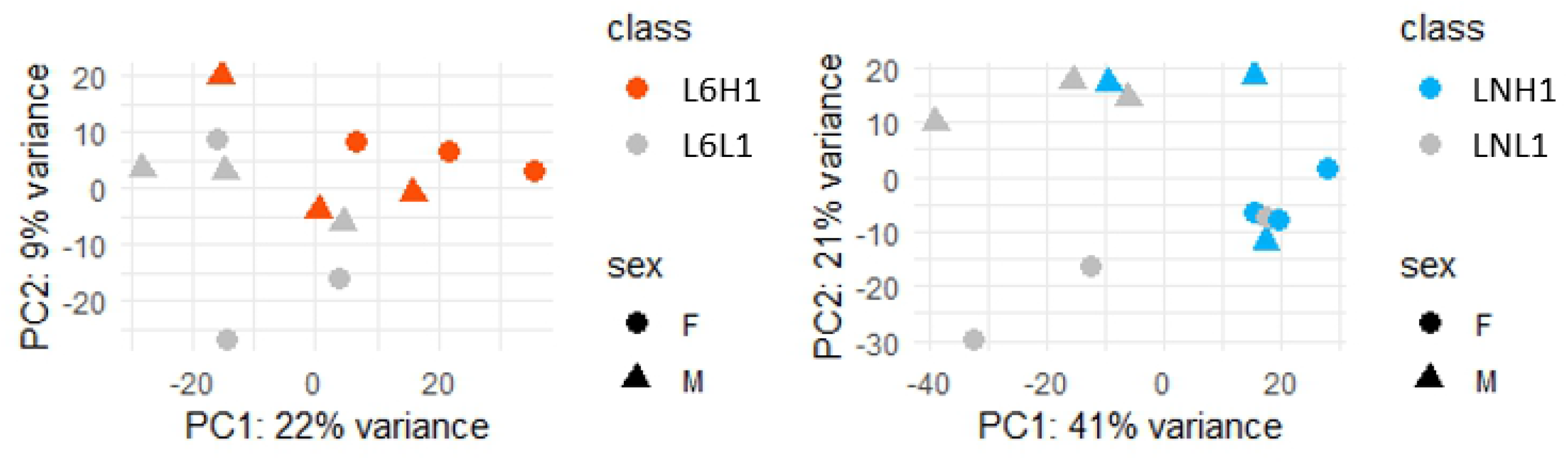
Principal component analysis of the gene expression in caecal tonsils from experiment 1 in each line. (A) Principal component analysis (PCA) of the gene expression in caecal tonsils within line 6_1_ in experiment 1. (B) PCA of the gene expression in caecal tonsils within line N in experiment 1.

A differential analysis with DESeq2 between low and high carriers within line 6_1_ in experiment 1 (L6L1/L6H1) allowed the identification of 97 DEGs (p-adj < 0.05), among which 42 were up- and 55 were down-regulated (S2 Fig and S2 Table). A similar analysis performed between low and high carriers within line N in experiment 1 (LNL1/LNH1) allowed the identification of 1,034 DEGs (p-adj< 0.05) with 794 up- and 240 down-regulated genes (S3 Fig and S3 Table). Only 1 DEG was shared between these two comparisons (Fig 4). In experiment 2, no significant DEGs were found between low and high carriers regardless of the line. The results are summarized in Fig 1. Merging both experiments in a single analysis, including a fixed effect for the experiment did not provide conclusive results.

**Fig 4.**
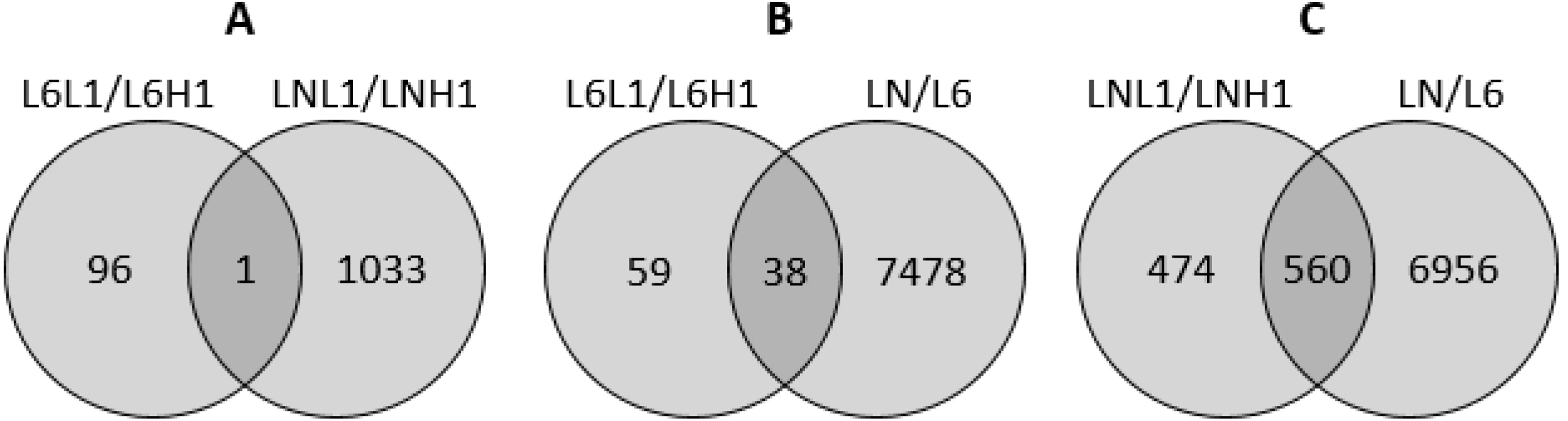
Venn diagram. Differentially expressed genes identified by comparing different groups of animals: (A) L6L1/L6H1 and LNL1/LNH1 (low vs high carriers of lines 6_1_ and N, respectively, in experiment 1), (B) L6L1/L6H1 and LN/L6 (low vs high carriers of line 6_1_ in experiment 1, and line N vs line 6_1_), and (C) LNL1/LNH1 and LN/L6 (low vs high carriers of line N in experiment 1, and line N vs line 6_1_).

### DEGs common to intra-line and between-line analyses

When comparing DEGs identified between lines 6_1_ and N, and within line 6_1_ between low and high carriers, 38 genes were shared. Among these shared genes, 9 genes appeared to be regulated in the same direction: 4 DEGs (*CEMIP, DMXL1, FUT2, NOS2*) were upregulated in both the resistant line N (low carriage) and in low carriers in line 6, and 5 DEGs (*PLEKHS1, RPS6KB2, CYP4B7, CYP2D6, CYP2AC1*) were downregulated in both line N and in low carriers in line 6_1_ (Fig 4 and S4 Table).

When comparing DEGs identified between lines 6_1_ and N and within line N between low and high carriers, 560 genes were shared. Among these shared genes, 58 appeared to be regulated in the same direction: 43 DEGs were upregulated in both the resistant line N (low carriage) and in low carriers in line N, and 15 DEGs were downregulated in both line N and in low carriers in line N (Fig 4 and S4 Table).

### Functional enrichment analysis

Enrichment analyses were performed with topGO on DEGs between all animals from lines N and 6_1_ and between low and high carriers within each line in experiment 1. The comparison between lines N and 6_1_ led to the identification of 62 significantly (p-value < 0.05) enriched Biological Processes (BP) Gene Ontology (GO) terms (S5 Table). The comparison between low and high carriers of lines 6_1_ (L6L1/L6H1) led to the identification of 23 BP GO terms (S6 Table). The comparison between low and high carriers of line N (LNH1/ LNL1) led to the identification of 288 BP GO terms (S7 Table).

The results are summarized in Fig 1. Three BP GO terms were shared between the L6L1/L6H1 and N/6 analyses (Table 1), fourteen between the LNL1/LNH1 and LN/L6 analyses (Table 2) and four between the L6L1/L6H1 and LNL1/LNH1 analyses (Table 3). Interestingly, the 3 GO BP terms enriched and shared between L6L1/L6H1 and N/6 were related to the response to biotic stimulus and other organisms.

**Table 1.** BP GO terms shared between LN/L6 and L6L1/L6H1 analyses.

| GO.ID | Term |
| --- | --- |
| GO:0009607 | response to biotic stimulus |
| GO:0043207 | response to external biotic stimulus |
| GO:0051707 | response to other organism |

**Table 2.** BP GO term shared between LN/L6 and LNL1/LNH1 analyses.

| GO.ID | Term |
| --- | --- |
| GO:0003413 | chondrocyte differentiation involved in endochondral bone morphogenesis |
| GO:0003416 | endochondral bone growth |
| GO:0003417 | growth plate cartilage development |
| GO:0003418 | growth plate cartilage chondrocyte differentiation |
| GO:0006644 | phospholipid metabolic process |
| GO:0030835 | negative regulation of actin filament depolymerization |
| GO:0030837 | negative regulation of actin filament polymerization |
| GO:0032272 | negative regulation of protein polymerization |
| GO:0043242 | negative regulation of protein complex disassembly |
| GO:0043244 | regulation of protein complex disassembly |
| GO:0051494 | negative regulation of cytoskeleton organization |
| GO:0051693 | actin filament capping |
| GO:0098868 | bone growth |
| GO:1901880 | negative regulation of protein depolymerization |

**Table 3.**
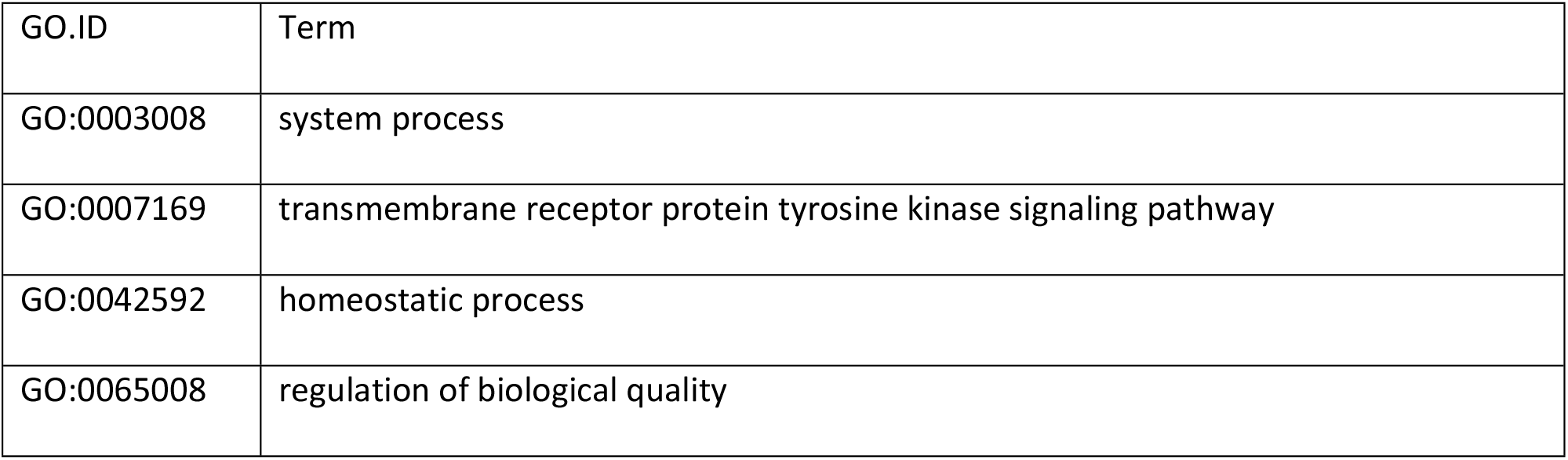
BP GO term shared between L6L1/L6H1 and LNL1/LNH1 analyses.

| GO.ID | Term |
| --- | --- |
| GO:0003008 | system process |
| GO:0007169 | transmembrane receptor protein tyrosine kinase signaling pathway |
| GO:0042592 | homeostatic process |
| GO:0065008 | regulation of biological quality |

## Discussion

We explored the caecal tonsil transcriptome after *Salmonella* Enteritidis infection by comparing samples from two genetic lines displaying contrasted levels of *Salmonella* carriage after infection. Differentially expressed genes were identified between lines N and 6_1_ and between low and high carriers within each line.

### A large difference of gene expression between lines

We showed a strong impact of the genetic background on gene expression in caecal tonsils, with 7,516 significant DEGs and 62 GO BP terms identified between the two lines. This strong difference in gene expression between lines is likely the result of genetic differences between the two lines. Some of these genes could explain the differences in susceptibility to *Salmonella* between lines. Thus, several genes display functions which may explain the higher resistance of line N, in which they were expressed to a greater extent. The latter genes code for the major histocompatibility complex I or II (MHCIBF2, MHCIYF5, MHCIIBLB1, MHCIIBLB2), antimicrobial peptides such as granzyme A and K, or the avian beta-defensines 10, 13 and 14. In the same way, the genes TLR_1,2,4,7,15_ [31–34], NOS2, Gal 13, PSAP and IGL, which have already been associated with SE resistance in a genetic study in chicken [13], were more expressed in the resistant line N.

Interestingly, we showed previously that the caecal microbiota composition of these animals was highly different between individuals from lines N and 6_1_ [30]. Do some of these DEGs in caecal tonsils indirectly impact *Salmonella* carriage through the modulation of the caecal microbiota composition? Conversely, do differences of microbiota composition indirectly impact *Salmonella* carriage through the modulation of gene expression in caecal tonsils? Further elements are needed to answer these questions.

### A difference of gene expression between low and high carriers within line

The comparisons between low and high carriers within each line in experiment 1 (L6L1/L6H1 and LNL1/LNH1) revealed only one DEG and four BP GO terms in common, leading us to the conclusion that host pathways leading to a higher resistance could differ between lines. It may be explained by their genetic differences. However, one of the four common BP GO terms may be directly related to the *Salmonella* infection process: the transmembrane receptor protein tyrosine kinase signalling pathway (GO:0007169), which is enriched in low carriers in both lines. Several studies have shown that the activation of protein tyrosine kinases and other specific transcription factors directly affects the innate immune response during SE infection [5,35]. Therefore, the transcriptional up-regulation of this pathway may be one of the mechanisms by which the animals from both lines control the infection.

Interestingly, in spite of the smaller difference between low and high carriers observed in line N compared to line 6_1_, we identified many more DEGs and BP GO terms in line N, compared to line 6_1_ (1034 DEGs and 288 BP GO terms in line N vs. 97 DEGs and 23 BP GO terms in line 6_1_). Previously, we showed that the microbiota composition was significantly different between low and high carriers in line 6_1_ but not between groups in line N [30]. One tentative hypothesis could be that the largest variability in *Salmonella* carriage in line 6_1_ could be explained by differences in gut microbiota composition, as demonstrated with the PA12 chicken line [24]. The resistance against *Salmonella* in line 6_1_ may thus be driven by mechanisms involving the intestinal microbiota, whereas in line N resistance mechanisms could be triggered by other factors, such as intra-line genetic variations.

Even in controlled environmental conditions with inbred lines, variability in the level of *Salmonella* carriage was identified within each line, as expected with the use of isolators preventing inter-individual recontamination [2]. This variability may derive from the residual genetic variability remaining within each line, or it may be caused by variations in the caecal microbiota composition, which is highly variable between lines and within line 6_1_ [24,30]. However, this variability was much higher in experiment 1, allowing the identification of more contrasted low and high classes than in experiment 2. This higher variability, and thus greater contrast in host response to *Salmonella* infection between low and high carriers, is probably the reason why significant DEGs and BP GO terms could be identified in experiment 1 but not experiment 2. Experimental conditions were highly controlled and similar between experiments but some unchecked factors, such as the environmental microbial exposure at hatch, may differ and explain the observed differences between the two.

### Identification of genes associated with the response to *Salmonella* infection

To highlight host genes that may be consistently associated with resistance to SE infection, we decided to focus on common DEGs between the intra-line and inter-line analyses. Indeed, genes more highly expressed in both the resistant line N and in low carriers in the susceptible line 6_1_, may be more likely to be associated with the resistance to SE infection. We further investigated if some of these genes have already been associated with the control of *Salmonella* infection in previous studies, supporting the reliability of our analysis.

Four genes were identified as more highly expressed in both the resistant line N and in low carriers of the susceptible line 6_1_ and are therefore associated with *Salmonella* resistance. Three of these genes have functions that are related to *Salmonella*: *NOS2, DMXL1* and *FUT2*. First, *NOS2* is a gene coding for the inducible nitric oxide synthase (iNOS) protein involved in macrophage inflammatory response [36]. Its function in the innate immunity against bacteria, viruses, fungi and parasites is well established, especially against *Salmonella* Typhimurium in mice models [37–42] and in *in vitro* models [43,44]. In chicken, a transcriptome analysis of caeca showed an increase of *NOS2* expression in the SE infected group [6], while genetic studies showed an association between *NOS2* gene alleles and spleen SE bacterial load [45,46]. An *in vitro* test showed the implication of iNOS between pathogen and macrophage cells during SE infection [47]. Finally, CNP (chitosan-nanoparticle) vaccination seems to increase *NOS2* expression and protect against SE infection in chicken [48]. Second, *DMXL1* is a gene involved in the phagosome acidification in macrophages. It seems to have an impact on innate immunity and macrophage bacteria killing after an activation by the TPL-2 kinase [49]. Finally, *FUT2* is a gene coding for the α-1,2-fucosyltransferase enzyme involved in the glycosylation profile of the gastrointestinal tract. In mice and human, it has been shown that a “non-secretor” individual heterozygous for a loss-of-function mutation is more susceptible to chronic intestinal diseases such as Crohn’s disease and to pathogen infection [50]. More specifically, it has been shown that “non secretor” mice (*Fut*^-/-^) show an increase of *Salmonella* Typhimurium in caecal tissue compared to wild-type mice [51,52]. Moreover, a *FUT2* polymorphism was associated with the human faecal microbiota composition and diversity [53], which could explain host-microbe interactions and susceptibility to infection [54]. Finally, it has been shown that *FUT2* was associated with the abundance of *Christensenellaceae* [55], a bacteria family we identified as associated with *Salmonella* Enteritidis resistance in the same animals [30]. Thus, *FUT2* may be a gene indirectly associated with SE resistance through the modulation of the microbiota composition, modulating the abundance of competitive bacteria against *Salmonella*.

Five genes were identified as less expressed both in the resistant line N and in low carriers of the susceptible line 6_1_ and may therefore be associated with *Salmonella* susceptibility. Among these genes, three belong to the cytochrome P450 family: *CYP4B7, CYP2D6* and *CYP2AC1*. CYP enzymes play a key role in metabolic processes in the intestine as the metabolism of xenobiotic substances. Metabolism of CYP enzymes is closely connected with infection, inflammation and intestinal microbiota in human [56,57].

Forty-two genes were identified as more expressed both in the resistant line N and in low carriers of the resistant line N and may therefore be associated with *Salmonella* resistance. Eight of these genes have functions of interest: *MHCIBF2, CLDN7, SIRT5, ENTPD1, SYT7, SLC22A23, S100B* and *MUC4*. The *MHCIBF2* (*Major histocompatibility complex class I antigen BF2*) gene is the predominant ligand of cytotoxic T lymphocytes [58]. It has been shown that a particular MHC I haplotype may contribute to control the response to SE infection in chicken [59,60]. The *CLDN7* (*claudin 7*) gene is involved in the formation of tight junctions between epithelial cells. It seems that a downregulation of CLDN7 by pathogen could facilitate translocation of invasive bacteria across the epithelium [61]. The *SIRT5 (sirtuin 5)* gene could have large impact on cellular homeostasis and is more expressed in colorectal cancer [62]. The *ENTPD1 (ectonucleoside triphosphate diphosphohydrolase 1* or *CD39*) gene has an impact on inflammatory bowel disease (IBD) and regulation of pro-inflammatory responses and pathogen colonization [63–65], as does the *SLC22A23* (*Solute Carriers family*) gene, which is associated with intestinal inflammation in human [66]. The *SYT7* (*synaptotagmin VII*) gene is associated with the control of cytotoxic granule fusion in lymphocytes, and mice lacking *syt7* have reduced ability to clear an infection [67]. The following two genes could be indirectly associated with SE resistance through the modulation of the microbiota composition. The *S100B* (*S100 calcium binding protein B*) gene codes for a signalling molecule which could be implicated in the communication mechanisms between microbiota and gut, and could explain differences between healthy and pathological microbiota in human [68]. Finally, a decrease of *MUC4* (mucin 4, cell surface associated) gene expression has been associated with ST infection in pigs [69], and genetic variants in this gene have been associated with the increase of gene expression relative to immune function and gut homoeostasis [70,71]. Mucins favour the establishment and the maintenance of a commensal microbiota, and form a protection barrier against pathogens. In mice, a study showed that mucins, including MUC4, may limit bacterial access to the epithelial surface in a *C*. rodentium infection [72]. In general, many studies in chicken showed the association of the expression of mucin genes as *MUC2* with SE infection [73–75].

Fifteen genes were identified as less expressed both in the resistant line N and in low carriers of the resistant line N and may therefore be associated with *Salmonella* susceptibility. Four of these genes have immune functions: L*ECT2, DEFB4A, LOC121107850* and *EXP*. The *LECT2 (leukocyte cell derived chemotaxin 2)* gene has an antibacterial function, is expressed in chicken heterophils, and increases in abundance in macrophage after SE infection [53]. It is also more expressed in vaccinated chicken again SE. The *DEFB4A (defensin beta 4A)* gene has an important role in innate immunity in mucosal tissues through its antimicrobial activity against various microorganisms. It has been shown that beta-defensin plays a role in immunoprotection against *Salmonella* Enteritidis in *in vitro* embryonic chicken cell model [54] and in the development of innate immunity in gastrointestinal tract of newly hatched chicks [55]. The *LOC121107850 (T-cell-interacting, activating receptor on myeloid cells protein 1-like)* gene codes for an activating receptor on myeloid cells protein 1-like, and the *EPX (eosinophil peroxidase)* gene codes for an enzyme in myeloid, which has a function in bacterial destruction [76].

Finally, some of the BP GO terms identified are associated with the response to SE infection. The three BP GO terms identified both between lines N and 6_1_ and between low and high carriers in line 6_1_ (LN/L6 and L6L1/L6H1) have interesting links to immunity: response to biotic stimulus, response to external biotic stimulus and response to other organisms (Table 1). Seven of the fourteen BP GO terms identified both between lines N and 6_1_ and between low and high carriers in line N (LN/L6 and LNL1/LNH1) are related to the regulation of the polymerisation or depolymerisation of proteins as actin (Table 2). It has been shown that a cytoskeletal actin rearrangement is induced by SE invasion in host cells [77]. Indeed, the actin cytoskeleton is targeted by *Salmonella* to promote its invasion, survival and growth in cells [78]. Genes implicated in these pathways may contribute to response to *Salmonella* infection and could be under genetic control.

## Conclusion

To conclude, the two experimental genetic lines N and 6_1_, displaying contrasted levels of *Salmonella* carriage after infection, showed a large difference of caecal tonsil gene expression associated with the outcome to SE infection. The comparison of resistant chicks (from line N or from low carriers within both lines) and susceptible chicks (from line 6_1_ or from high carriers within both lines) allowed us to identify several genes and pathways associated with *Salmonella* resistance. Different mechanisms seem to be involved in the response to SE between these two experimental lines. A lower number of DEGs is associated with a larger inter-individual variability in line 6_1_ compared to line N. In light of our previous results obtained on the caecal microbiota composition of the same animals, the difference of microbiota composition between low and high carriers in line 6_1_ may be one of the explanations of this high intra-line variability. These results suggest that some of the genes identified, such as *FUT2*, could act indirectly on *Salmonella* carriage through the modulation of the microbiota composition.

## Materials and methods

### Experimental design

All animal procedures were authorised by the Ethic committee: APAFIS#5833-2016062416362298v3. Animals from the two experimental White Leghorn inbred lines N and 6_1_ were provided by the experimental unit PEAT (Pole d’Expérimentation Avicole de Tours, Nouzilly, France). As described previously [30], animals were raised together on the floor until infection at the PFIE unit (Plateforme d’Infectiologie Expérimentale, INRA, Nouzilly, France) with free access to food and water. At 7 days of age, chicks were orally infected with *Salmonella* enterica Enteritidis (Strain 775 [LA5 Nal20Sm500], 5.10^4^ cfu/0.2 mL/chick) and immediately separated into isolators. Four isolators were used for each experiment: two isolators for chicks from line N and two others for chicks from line 6_1_, with 30 birds per isolator. Caecal contents and caecal tonsils were collected at 12 days post infection after the animal sacrifice and were immediately frozen in liquid nitrogen and stored at -80°C until use. Two experiments were conducted with a total number of 240 chicks.

### DNA extraction, *Salmonella* count by Droplet Digital PCR and choice of low and high carriers

As described previously [30], individual caecal DNA was extracted from an average of 200 mg of frozen caecal contents and DNA samples were stored at -20°C. Individual abundances of *Salmonella* Enteritidis in caecal contents were obtained by Droplet Digital PCR (ddPCR) using the QX200 Droplet Digital PCR system (Bio-Rad) at the @bridge platform (INRAE, Jouy-en-Josas, France). Data were analysed with a log transformation of the copies of *Salmonella*. Analyses of variance (ANOVA) were performed to test the significance of differences of the copies of *Salmonella* according to different factors (line, sex, experiment or isolator) using the anova function from base R (Type I sum of squares). We selected 48 chicks, either low or high *Salmonella* carriers based on data from ddPCR, balancing the experiment, sex, isolator and line factors.

### RNA extraction and sequencing

Caecal tonsils from the 48 low/high carrier chicks were first grinded using ULTRA-TURRAX T25 (IKA). RNAs were then extracted using the NucleoSpin RNA Kit (MACHEREY-NAGEL) according to the manufacturer’s protocol. The quantity of RNA was measured using a Nanodrop spectrophotometer (Thermo Scientific) and its quality was assessed using a 2100 Bioanalyzer Expert system using a total RNA nano Kit (Agilent), all samples displayed an RNA integrity number (RIN) > 7. RNAs were sent to the genomic platform GeT-Plage (Toulouse, France) for the cDNA library preparation (TruSeq Stranded mRNA kit, Illumina) and the sequencing (NovaSeq 6000 Sequencing System, Illumina). The sequencing data analysed during the current study are available in the NCBI Sequence Read Archive (SRA) database under the Bioproject accession number PRJNA649900.

### Transcriptome analysis

The quality control of raw reads was performed using the FastQC program. Trimming was performed using Sickle (version 1.33). Reads were then mapped on the *Gallus gallus* reference genome (Gallus_gallus.GRCg6a.98.gtf) with STAR (version 2.5.3a) [79]. Genes were counted with htseq-count (version 0.12.4) [80].

The metadata and table of gene counts have been included as Additional files 6 and 7, respectively. PCA and ANOVA tests on PCA components were performed with the stats package (3.6.1). DESeq2 (version 1.26.0) was used with R version 3.6.1 to perform differential gene expression analyses [81]. In our model, the sex and the isolator effects were fixed. Genes with an adjusted p-value < 0.05 were considered to be differentially expressed genes (DEGs). The ggplot2 packages (3.2.1) were used for visualisation. The functional enrichment analyses were performed with topGO package version 2.38.1 [82].

## Acknowledgments

We thank our colleagues from the experimental unit PEAT who raised the parents of the animals studied and provided the chicks used in our study. We also thank colleagues from the experimental unit PFIE, who efficiently monitored the experiments and collected the samples, and colleagues from the SPVB research team, who helped collect the samples.

## Supporting information

**S1 File. ANOVA analysis results on PCA1 and PCA2 components S2 File. Metadata associated with all samples**

**S3 File. Gene count table**

**S1 Fig. DESeq2 results for the comparison of the N and 6**_**1**_ **lines**. Distribution of raw p-values (A), dispersion plot (B), and MA plot highlighting 1492 and 1285 up- and down-regulated genes with a p-adj < 0.05 (C).

**S2 Fig. DESeq2 results for the comparison of the low and high carriers within line 6**_**1**_ **in experiment 1**. Distribution of raw p-values (A), dispersion plot (B), and MA plot highlighting 42 and 55 up- and down-regulated genes with a p-adj < 0.05 (C).

**S3 Fig. DESeq2 results for the comparison of the low and high carriers within line N**. Distribution of raw p-values (A), dispersion plot (B), and MA plot highlighting 794 and 240 up- and down-regulated genes with a p-adj < 0.05 (C).

**S1 Table. Differentially expressed genes between LN and L6 groups**

**S2 Table. Differentially expressed genes between L6H1 and L6L1 groups S3 Table. Differentially expressed genes between LNL1 and LNH1 groups**

**S4 Table. Common differentially expressed genes between LN/L6 analysis and L6L1/L6H1 analysis and between LN/L6 analysis and LNL1/LNH1 analysis**

**S5 Table. Enrichment analyse results performed on DEGs between line N and 6**_**1**_
**S6 Table. Enrichment analyse results performed on DEGs between low and high carriers in line 6**_**1**_ **-experiment 1**

**S7 Table. Enrichment analyse results performed on DEGs between low and high carriers in line N - experiment 1**

